# Digital dissection of arsenate reductase enzyme from an arsenic hyperccumulating fern *Pteris vittata*

**DOI:** 10.1101/056036

**Authors:** Zarrin Basharat, Deeba Noreen Baig, Azra Yasmin

## Abstract

Action of arsenate reductase is crucial for the survival of an organism in arsenic polluted area. *Pteris vittata*, also known as Chinese ladder brake, was the first identified arsenic hyperaccumulating fern with the capability to convert [As(V)] to arsenite [As(III)]. This study aims at sequence analysis of the most important protein of the arsenic reduction mechanism in this specie. Phosphorylation potential of the protein along with possible interplay of phosphorylation with *O*-β-GlcNAcylation was predicted using neural network based webservers. Secondary and tertiary structure of arsenate reductase was then analysed. Active site region of the protein comprised a rhodanese-like domain. Cursory dynamics simulation revealed that folds remained conserved in the rhodanese main but variations were observed in the structure in other regions. This information sheds light on the various characteristics of the protein and may be useful to enzymologists working on the improvement of its traits for arsenic reduction.

## Introduction

Arsenic contamination in soil and groundwater poses significant threat to human health and cleanup of arsenic contaminated soil and water using phytoremediation approach is a cost effective strategy (Meadows, 2014). Presence of arsenic in the environment has led to the adaptation of various prokaryotic and eukaryotic species with a repertoire of enzymes for detoxification of this metalloid and hence, aid cleanup of the environment. The initial and crucial step of arsenic reduction involves conversion of arsenate [As(V)] to arsenite [As(III]). This is accomplished by arsente reductase enzyme which catalyzes reduction of [As(V)] to [As(III]) and vice versa (Martin et al., 2001). Arsenate reductase belongs to the family of oxidoreductase enzymes, especially the ones that act with phosphorus or arsenic being donor and disulfide acting as acceptor. Similarity of arsenate to phosphate makes it perfect for ready plant uptake of arsenate as it is mistaken for plant nutrient group phosphate (Ullrich-Eberius et al., 1989; Meharg and McNair, 1992; Dhankher et al., 2006).

*Pteris vittata,* also known as the Chinese brake or Ladder brake, is an arsenic hyperaccumulating fern and used as a model in arsenic pollution mitigation studies (Wilkins and Salter, 2003). Root uptake of [As(V)] is followed by reduction to [As(III)]. [As(III)] is then transported to the lamina of the *Pteris vittata* frond and deposited in the cells as free arsenite (Indriolo et al., 2010). The frond can horde up to 22,630 mg As kg^−1^ (Wang, 2002). *Physcomitrella patens* is a moss specie and belongs to one of the oldest species of plants that inhabited land (Heckman, 2001). Sequence, topology, structure and functional information is critical for the master regulator of arsenic hyperaccumulation in this plant as the expression, pathway and pollutant homeostasis network associated with it, that supports hyperaccumulation or tolerance can be important for environmental cleanup (Basharat and Yasmin, 2016a). This rationale of this study was to analyse arsenate reductase in this hyperaccumulating specie, using informatics tools, which is a quick and handy method of studying protein of interest.

## Material and Methods

Sequence comprising of 435 amino acid residues of arsenate reductase for *Pteris vittata* was retrieved from NCBI database with accession no: ABD64881. ProtParam tool(http://web.expasy.org/protparam) was used for physiochemical property analysis of the protein, while subcellular localization was calculated using CELLO v.2.5 (http://cello.life.nctu.edu.tw/) as previously described (Basharat, 2015). Conserved domains were studied using Conserved domain database (Marchler-Bauer et al., 2012). Position of phosphorylation sites in arsenate reductase of both species was studied using neural network based tool NetPhos 2.0 (Blom et al., 1999; Basharat and Yasmin, 2016b; Basharat and Yasmin, 2016c). It makes predictions for possible phosphorylation potential at serine, threonine and tyrosine sites in the eukaryotic proteins. Kinases responsible for phosphorylation were predicted using another neural network based tool NetPhosK (Blom et al., 2004). Possible serine or threonine sites that could act as substrates for phosphate as well as showed potential for *O*–ß–GlcNAcylation were predicted using YinOYang server (Gupta and Brunak, 2001).

1D structure was studied using Protter (http://wlab.ethz.ch/protter). Secondary structure motifs, wiring diagram of the protein was obtained by PROMOTIF program (Hutchison and Thronton, 1996) implemented at PDBSum server web interface (Laskowski, 2001; Laskowski et al., 1997; Laskowski, 2004; Laskowski, 2009). The tertiary structure of arsenate reductase was modeled using I-TASSER (Zhang, 2004; Roy et al., 2010; Yang et al., 2014). Models were built on the template structures of proteins from PDB library, identified by LOMETS (Wu and Zhang, 2007) and ranked as best template by threading programs in I-TASSER. Final model was chosen based on confidence or C-score, centred on the comparative clustering of structural density as well as consensus significance score of multiple threading templates. Structure quality assessment was done by Ramachandran plot analysis (Ramachandran et al., 1963; Morris et al., 1992; Lovell et al., 2003). Percentage of residues in favourable regions was calculated. ERRAT webserver (Colovos and Yeates, 1993) was also used to assess quality of structure. Model was rendered in Pymol v1.4.1. Missing hydrogens were added and energy minimization was performed with Molecular operating Environment (MOE). Dynamics simulation was carried out for 500 ps using MOE. GROMOS forcefield was applied, with NVT statistical ensemble for conformation generation and Nose-Poincare-Anderson algorithm (Bond, 1999) to solve the equation of motion. Timestep of 2 fs was selected for discretizing the equation of motion. Temperature (300K) and pressure (101 KPa) remained constant.

## Results and Discussion

In plants, few detailed studies have been conducted on arsente reductase enzyme. Arsenate reductase has been reported in *Arabidopsis thaliana* (Chao et al., 2014; Sánchez-Bermejo et al., 2014), *Oryza sativa* (Duan et al., 2007), green alga *Chlamydomonas reinhardtii* (Yin et al., 2011) and the arsenic hyperaccumulator *Pteris vittata* (Duan et al., 2011; Ellis et al., 2006; Cesaro et al., 2015) till to date. Here, *in silico* analysis has been attempted for this enzyme at sequence and structure level. Molecular weight of the protein was determined as 45271.7 daltons, theoretical pi: 9.31, total number of negatively charged residues (Asp + Glu): 37, total number of positively charged residues (Arg + Lys): 46, instability index (II): 29.98 (which classified the protein as stable). Subcellular localization analysis based on support vector machine technique revealed that the protein resided in the chloroplast region.

Protein phosphorylation represents the addition of a phosphate group to the amino acid side chain of an amino acid by a protein kinase (Tan, 2011). A phosphorylated amino acid may be dephosphorylated by a phosphatase or replaced by some other post translational modification as *O*–β–GlcNAcylation (Basharat and Yasmin, 2015). Phosphorylation can modulate the enzymatic activity, so it is necessary to study phosphorylation profile for gaining insights into protein function. A total of 34 phosphosites (19 serine, 10 threonine, 5 tyrosine) were predicted in *Pteris vittata* arsenate reductase while interplay of phoshosites wih *O-$-* glycosylation was observed at six sites (Table 1). Conserved domain analysis revealed that a rhodanese-like homology domain was present in the sequence at position 262–376 with an E–08 value threshold of 1.00 e^−08^. Rhodanese-like domain is an alpha beta fold domain with a cysteine residue comprising active site, characteristic of stress proteins as cyanide and arsenate resistance proteins (Bordo, 2002).

**Table 1.** Phosphorylation and *O*–β–GlcNAcylation potential of arsenate reductases in *Pteris vittata*. – indicates a complete absence or lack of information for specified residues. + indicates presence of a particular trait in the enzyme at mentioned site.

| **Residue** | **No.** | **Kinases**<br>**responsible for**<br>**Phosphorylation** | ***O*-β-**<br>**GlcNAcylation** |
| --- | --- | --- | --- |
| Ser | 12 | - | + |
|  | 26 | - | - |
|  | 27 | PKA | - |
|  | 29 | - | - |
|  | 52 | - | - |
|  | 65 | - | - |
|  | 85 | - | + |
|  | 86 | - | + |
| Thr | 89 | p38MAPK, cdk5 | - |
|  | 95 | - | + |
|  | 166 | - | + |
|  | 213 | CKII | + |
|  | 217 | CKI | - |
|  | 245 | - | - |
|  | 286 | - | - |
|  | 290 | PKC | - |
|  | 318 | - | - |
|  | 358 | - | - |
|  | 399 | - | - |
|  | 110 | - | - |
|  | 123 | - | - |
|  | 134 | - | - |
|  | 143 | - | - |
|  | 188 | - | - |
|  | 200 | - | - |
|  | 279 | - | - |
|  | 408 | - | - |
|  | 415 | - | - |
|  | 423 | - | - |
| Tyr | 19 | - | - |
|  | 145 | - | - |
|  | 249 | INSR | - |
|  | 272 | - | - |
|  | 299 | INSR | - |

1D plot revealed that the protein has a transmembrane region (residue 224–249) (Fig. 1). Secondary structure analysis disclosed presence of 1 sheet, 1 beta hairpin, 3 strands, 26 helices, 53 helix-helix interacs, 43 beta turns and 9 gamma turns (Fig. 2). 3D structure (Fig. 3) quality analysis using Ramachandran plot analysis revealed that 85.0% residues existed in the favoured, 10.2% in allowed regions while 4.8% as outliers. ERRAT displayed a value of 94.84 (~95%). This indicated the good quality of structure as good resolution structures have an ERRAT value of around 95% or higher. Short timescale dynamics simulation was carried out to study initial changes in protein structure conformation. Simulation analysis showed Cα (protein backbone) fluctuations and large variation in protein conformation was observed (Fig. 4). Larger timescale dynamics simulation is further required to study protein behaviour and map fluctuations and reoccurrence of native state accurately.

**Fig. 1.**
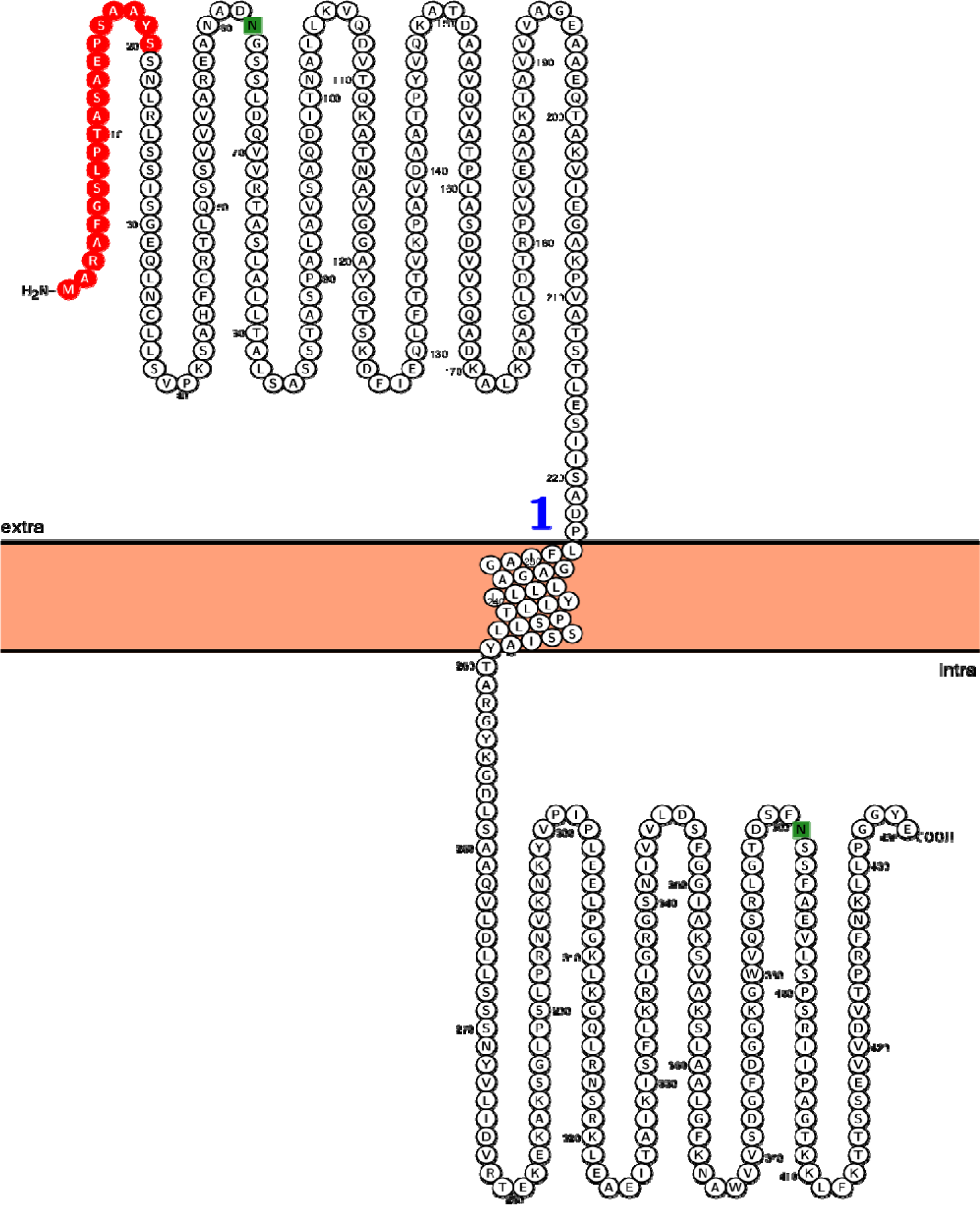
Topology of the arsenate reductase protein. Red colour shows the signal peptide region. N-glyco motif is shown in green colour.

**Fig. 2.**
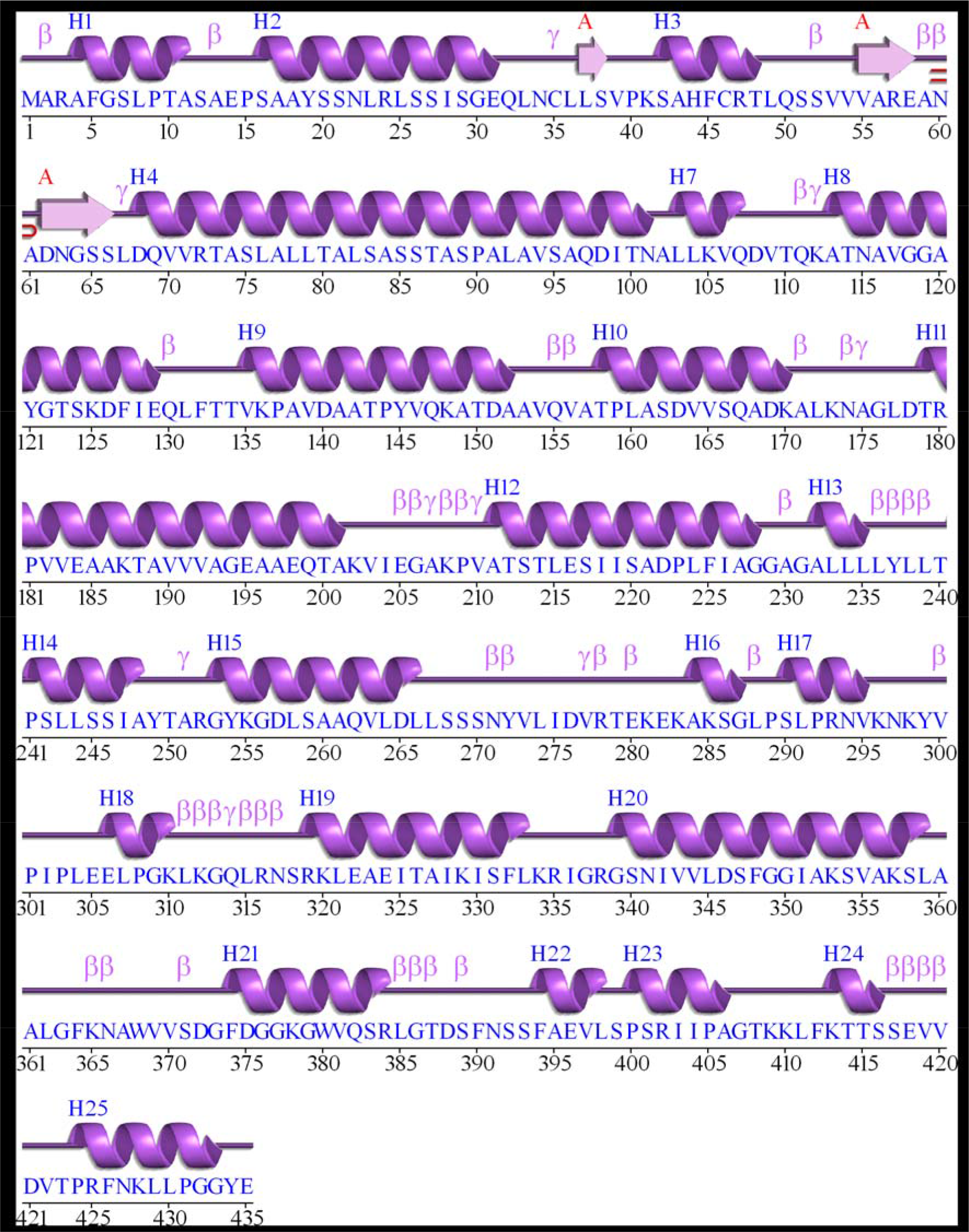
Schematic secondary structure wiring diagram of arsenate reductase for *Pteris vitatta*. Helices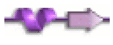 are labelled as H1, H2,… and strands by their sheets A and B. Motifs β and γ turns are also labelled.

**Fig. 3.**
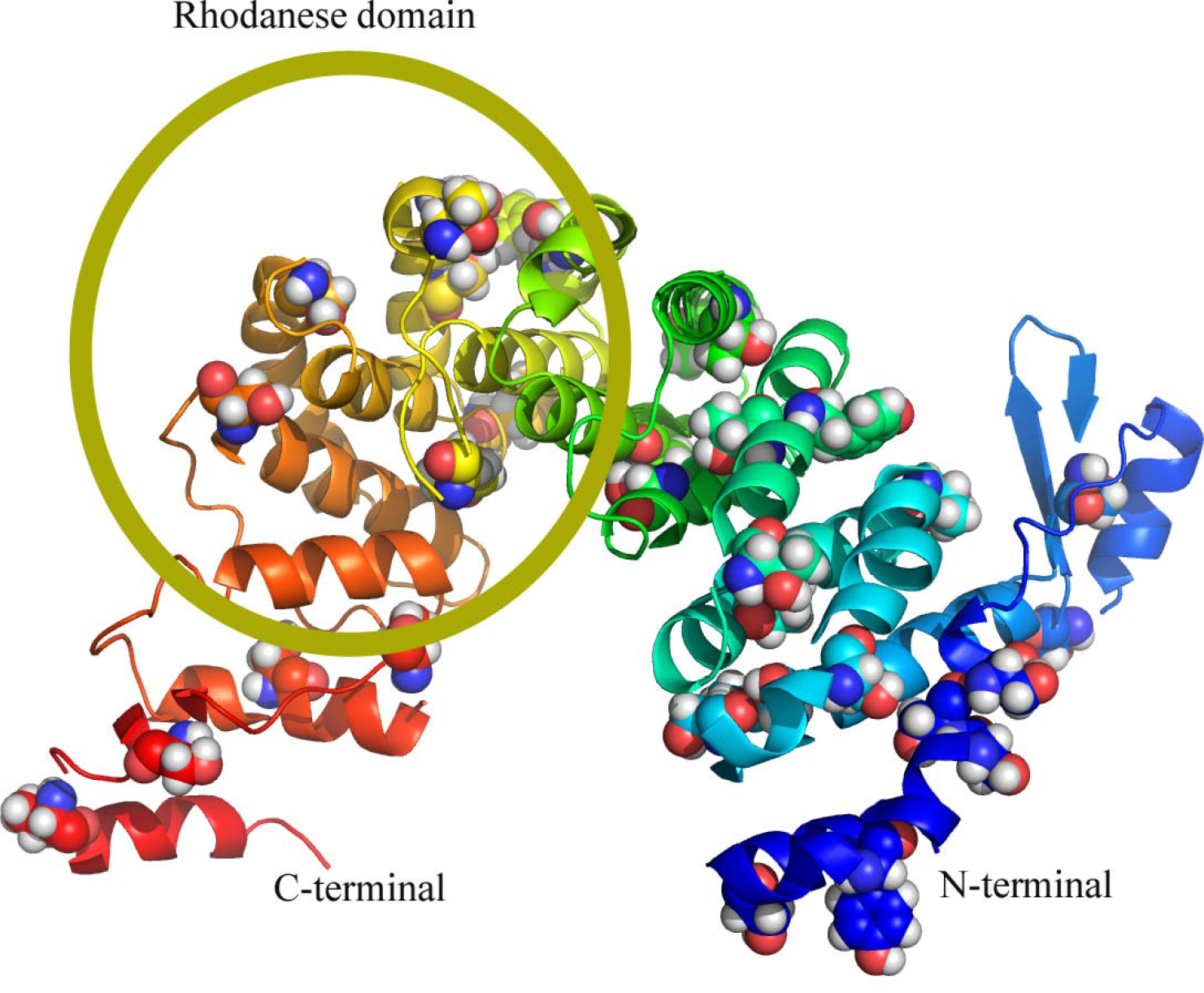
Modeled 3D structure of *Pteris vittata*. C-terminal is depicted by red and N-terminus by blue colour. Phosphorylated sites are depicted by spheres while rhodanese-like domain is encircled.

**Fig. 4.**
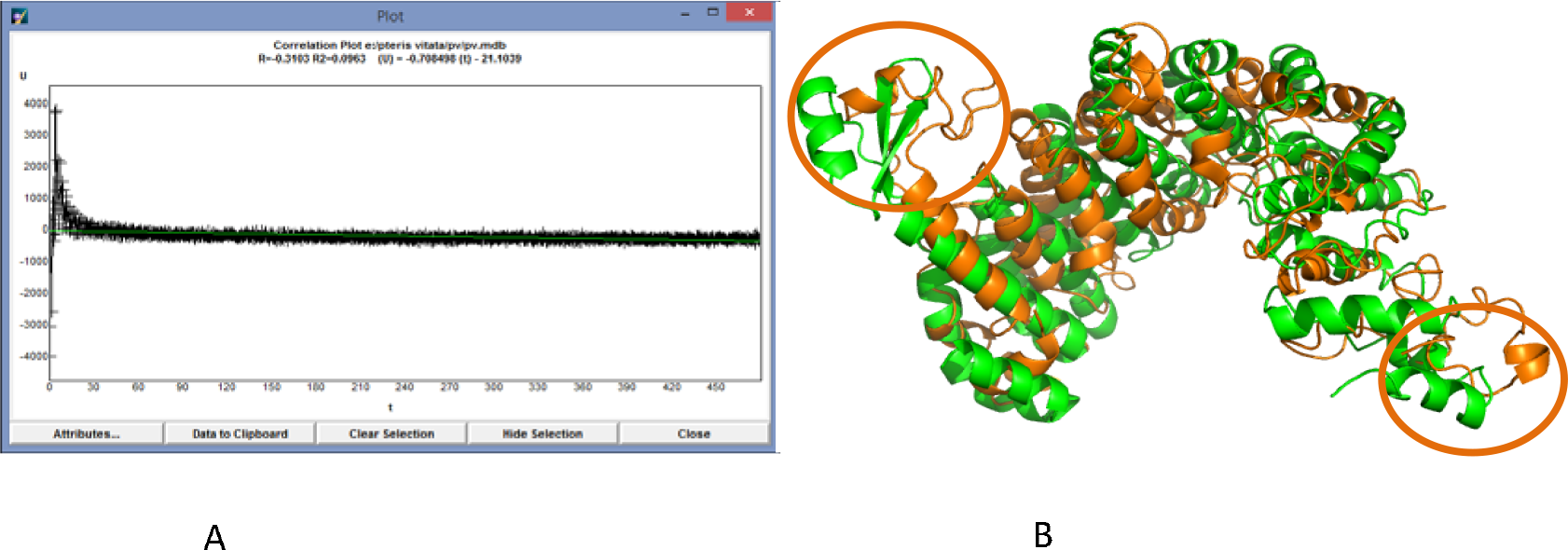
(A) Correlation plot of energy of the simulated protein versus time. (B) Cα fluctuation of native (green) and simulated (green) protein state for 500 ps. Beta sheets (residue 55–58 and 61–65) conform to loops. Helices in the rhodanes domain remained stable whereas helix at the C-terminal lost conformation to form loop (encircled).

This study is a useful addition to arsenate reductase knowledgebase and has shows that bioinformatics analysis of a protein sequence is a quick method for inferring functional properties. It also paves way for investing future research efforts focusing on comparative analysis of close and distant homologs as a wealth of information for this enzyme lies unexplored in other plants that have recently been sequenced. Evolutionary dynamics of protein phosphorylation occurring at the residue level should also be attempted. Analysis of protein-protein interactions with arsenate reductase helper enzymes is currently being conducted by the authors, which can further enhance the understanding of evolution and its impact in this enzyme system in closely related species. Experimental research in this domain is proposed and would definitely lead to new post translational modification prospects in arsenic reduction phenomenon in hyperaccumlators, as well as other plants.

